# Functional and taxonomic comparison of mouse and human gut microbiotas using extensive culturing and metagenomics

**DOI:** 10.1101/2021.02.11.430759

**Authors:** Benjamin S. Beresford-Jones, Samuel C. Forster, Mark D. Stares, George Notley, Elisa Viciani, Hilary P. Browne, Nitin Kumar, Kevin Vervier, Alexandre Almeida, Trevor D. Lawley, Virginia A. Pedicord

**Affiliations:** Cambridge Institute of Therapeutic Immunology and Infectious Disease, Jeffrey Cheah Biomedical Centre, Cambridge Biomedical Campus, Cambridge, UK, CB2 0AW; Department of Medicine, University of Cambridge School of Clinical Medicine, Cambridge Biomedical Campus, Cambridge, UK, CB2 2QQ; Wellcome Sanger Institute, Wellcome Genome Campus, Hinxton, UK; European Bioinformatics Institute (EMBL-EBI), Wellcome Genome Campus, Hinxton, UK

## Abstract

Mouse models are essential for biomedical science and drug discovery, yet it is not known how the bacteria in the mouse microbiota – important determinants of phenotypes of health and disease –affect their relevance to human disease. To interrogate the taxonomic and functional differences between the human and mouse gut microbiotas, we developed the Mouse Microbial Genome Collection (MMGC), a compilation of 276 genomes from cultured isolates and 45,218 metagenome-assembled genomes (MAGs) from 1,960 publicly available mouse metagenomes. The MMGC reveals that while only 2.65% of bacterial species are shared between mouse and human, over 80% of annotatable functions are present in both microbiomes. Using drug metabolism and butyrate synthesis as examples, we illustrate that although the species harbouring these key functions can differ between hosts, the MMGC enables identification of functionally equivalent taxa in the mouse and human microbiotas. The MMGC thereby facilitates the informed use of mice in biomedical research by providing access to the conservation and taxonomic locations of bacterial functions of interest.

## Introduction

Mouse models are essential tools for controlled experimental studies of conserved aspects of human physiology and disease. However, while the importance of the microbiota in determining phenotypes of health and disease of the host is well established^1-4^, relatively little is known about the mouse microbiome and therefore its impact on the use of mice as models for human biology. Prior to 2020, there existed fewer than 200 published mouse-derived gut commensal genomes, compared to more than 200,000 from human hosts^5^. Recent studies have started to bridge this gap^6-8^; however, efforts remain small-scale and non-systematic, and the difference in number of human and mouse commensal genomes remains stark. Recent studies have estimated a taxonomic overlap of 4-5% between human and mouse hosts^9, 10^, but it is unknown which species are shared and whether these estimates are accurate at the genome level. Comparing functional annotations of human and mouse metagenomes has yielded estimates of up to 95% functional crossover^9^; however, the taxonomic locations of these functions are not known nor their implications for medical and pharmaceutical research.

Both culture-dependent and -independent techniques have been successfully implemented to characterise the human microbiota^11-16^. The former involves isolating bacteria from faecal or luminal samples by high-yield culturing^11-13^ or by utilising a reverse genomics approach^14^, followed by whole genome sequencing. Culture-independent methods most often assemble shotgun metagenomes *de novo* and bin contigs into potential genomes called metagenome-assembled genomes (MAGs)^7, 12, 15, 16^. A combination of these methods has allowed for the construction of human microbiota genome and culture collections^5^ that improve metagenomic analyses^11^ and facilitate experimental validation of resulting correlative associations using cultured representatives^17, 18^. No such comprehensive collection currently exists for the mouse microbiota.

Here, we use an integrative approach, complementing high-throughput mouse gut commensal culturing with large-scale MAG synthesis from published mouse metagenomes, to establish the Mouse Microbial Genome Collection (MMGC), a comprehensive genome and culture repository. The MMGC contains 18,075 non-redundant, near-complete mouse-derived gut bacterial genomes (1,021 species; 766 novel) that significantly expand our understanding of the global mouse microbiome. It includes 276 whole genome-sequenced mouse commensal bacterial isolates that we have deposited in a public repository, increasing the number of isolate genomes previously published in a single study by 219%, making it the largest and most comprehensive collection of its kind to date. Leveraging the MMGC, we establish the taxonomic and functional overlap between human and mouse microbiotas and identify functionally equivalent species in each host. Utilising examples of drug metabolism and butyrate synthesis by the microbiota, we further illustrate how host-specific differences in the presence and location of these functions may have implications for pharmaceutical and biomedical research. This study provides an unprecedented understanding of human-mouse microbiota similarities and differences, facilitating the informed use of mouse models for the investigation of microbiota-susceptible phenotypes.

## Results

### A comprehensive collection of mouse-derived gut microbial genomes

To address some of the current limitations to the analysis of mouse metagenomic data, we performed high-throughput culturing of faeces from conventionally-housed laboratory mice alongside metagenome assembly and binning of publicly available mouse metagenomes.

From 30 specific-pathogen free (SPF) mice, we cultured 276 bacterial strains representing 130 species, including 73 that could not be assigned to a species within the Genome Taxonomy Database (GTDB), termed “novel” species^19, 20^ (Figure S1a, Supplementary Table 1). For generating MAGs, we used 1,960 publicly available SPF mouse gut metagenomes, of which 1,927 passed quality control, representing 71 studies and 19 strains of mice from 56 institutes across 18 countries and four continents (Supplementary Table 2). Only samples that had not received treatments that could disrupt the microbiota, such as antibiotics, were included. This allowed us to build a global collection, henceforth referred to as the MMGC, of isolate genomes and near-complete MAGs representing a total of 15 phyla and 1,021 species from the mouse intestinal microbiota, of which 766 (75%) were previously uncharacterised (Figure 1a). Of these 1,021 species, 12.7% have a representative isolate in our Mouse Culture Collection (MCC), allowing for experimental follow-up of the correlative inferences generated in silico^17^.

**Figure 1.**
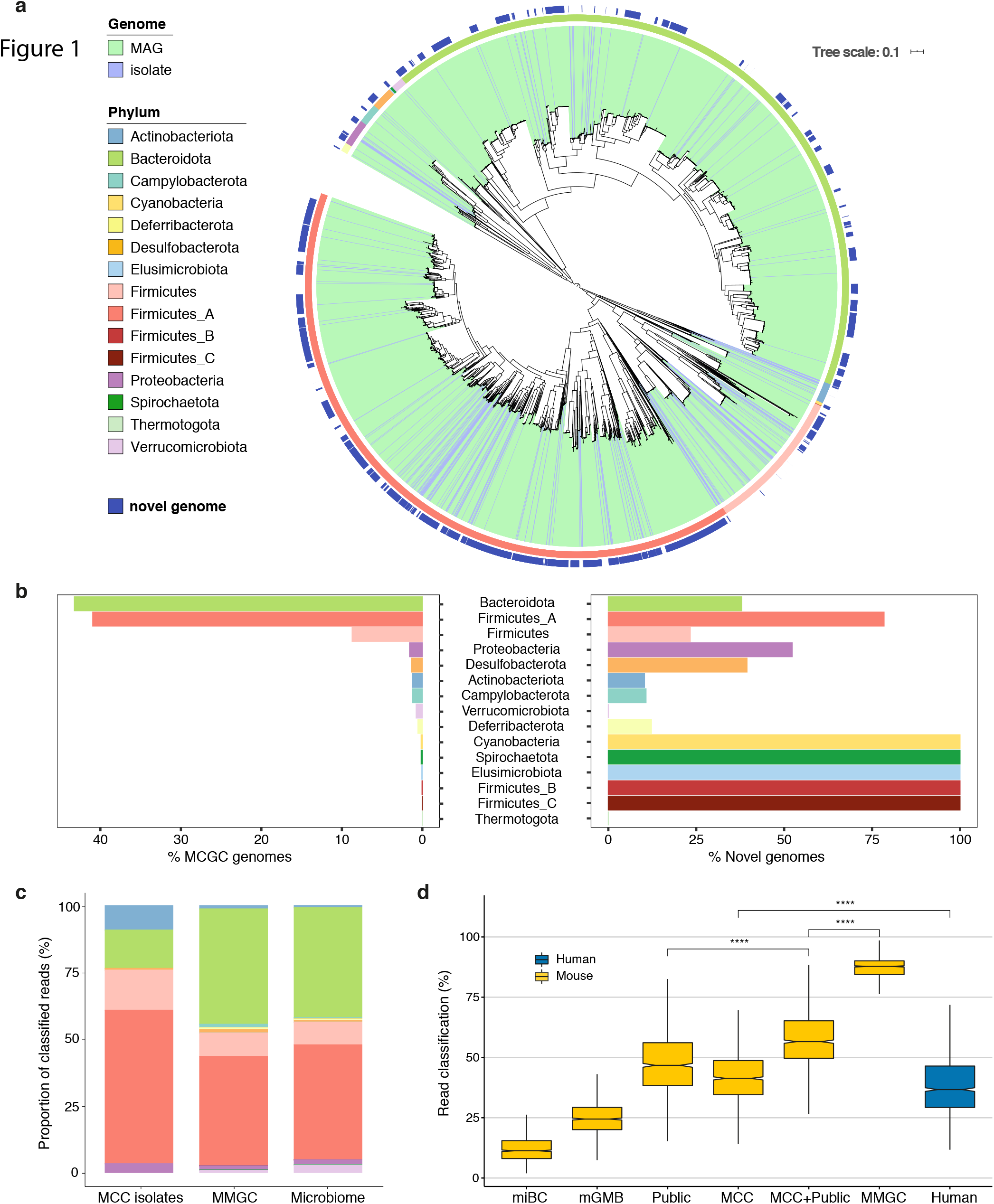
The MMGC is highly representative of the mouse microbiome. a) Maximum-likelihood tree of the 18,075 near-complete genomes of the MMGC, generated by aligning 120 core bacterial markers. Colour range represents genome-type, the inner colour bar denotes genome phylum, and the outer colour bar indicates whether the genome could not be assigned to a species taxonomic rank (“novel genome”). b) Distribution of MMGC genomes across phyla (left) and percentage of novel genomes by phylum (right). c) Phylum distribution of the 276 MMGC isolates (MCC isolates), all 18,075 MMGC genomes (MMGC), and the average mouse microbiome (Microbiome; n=1924). The distributions of each stacked bar were compared using a Chi-Square Test for Independence, MCC:Microbiome (*P* = 0.013, significantly different), MMGC:Microbiome (*P* = 0.999, not significantly different). d) Read classification rates of 1,259 mouse metagenome samples using different custom Kraken2 databases. The Human database contains near-complete representative genomes from the UHGG. miBC, n=43; mGMB, n=100; Public (miBC, mGMB, NCBI mouse isolates), n=274; MCC, n=276; MCC+Public, n=550; MMGC, n=1,021; Human, n=3,006. Significance was determined for selected comparisons using paired t-tests, \**P* <<<< 0.0001.

Using single-sample assembly and binning we generated 45,218 MAGs, all of which exceeded published criteria for medium quality^21^. Of these MAGs, 23,064 were sufficiently high quality to be defined as near-complete^22^ (Figure S1b, Supplementary Table 3). Most MAGs pertained to the classes Clostridia or Bacteroidia (Figure S1c). Using only near-complete genomes in our analyses, we combined our isolate genomes with our MAGs and removed highly related genomes (≤0.001) Mash distance^23^, equivalent to 99.9% ANI) to produce a non-redundant database of 18,075 genomes genomes (Supplementary Table 4). We next examined novel characteristics of the mouse microbiota revealed by the MMGC. Bacteroidota and Firmicutes_A were the most dominant taxonomic phyla represented with 43.2% and 40.9% of genomes, respectively (Figure 1b). Firmicutes_A dominated the novelty in the MMGC, with 78.4% or 5,801 genomes previously uncharacterised (Figure 1b). Phyla such as the Cyanobacteria and Spirochaetota, while relatively minor members of the collection, were represented entirely by novel species. The MMGC is highly representative of the average mouse microbiota (Figure 1c; *P* = 0.999, Chi-Square Test of Independence), demonstrating the utility of our complementary approach in assaying both culturable and currently uncultured fractions of the host microbiota.

To ascertain whether the MMGC improves taxonomic classification of mouse metagenomes, we compared read classification rates of samples using different collections of mouse commensal genomes^6, 8^ and all public mouse commensal genomes available on NCBI (Figure 1d). Using only the representative genomes for each species in the MMGC, and analysing 1,259 independent metagenomic samples, the MMGC on average classified 87.7% of reads, compared to 46.3% with public genomes alone. The MMGC significantly increased read classification in comparison to all other databases, and therefore offers previously unattainable coverage of the mouse microbiome. Of note, a representative human gut microbial database, the Unified Human Gastrointestinal Genome (UHGG) collection, was comparably poor for analysis of mouse metagenomes, classifying only 36.6 % of reads (Figure 1d). Together, our culture-dependent and -independent approaches generated a highly representative collection of cultured isolates and a database of mouse gut bacterial genomes that enable both significantly improved interpretation of mouse metagenomic data and experimental follow-up in mouse models.

### Population structures of the mouse gut microbiota

As the MMGC offered an unprecedented ability to analyse the mouse microbiota, we next sought to accurately define the average mouse microbiome by taxonomic classification at species-level resolution. We used the MMGC to analyse 1,785 publicly available, globally distributed mouse shotgun metagenomes, including samples from both laboratory and wild mice (Supplementary Table 5). Species of phylum Firmicutes_A represented 18 of the top 20 most prevalent species in the mouse microbiota (Figure 2a); however, they only represented two of the top 20 most abundant species (Figure S2). In contrast, members of phylum Bacteroidota were much more abundant, with 15 representatives in the top 20 most abundant species (Figure S2a). *CAG-485 sp002362485*, a member of the Muribaculaceae family (p. Bacteroidota), was the most abundant species while also being highly prevalent.

**Figure 2.**
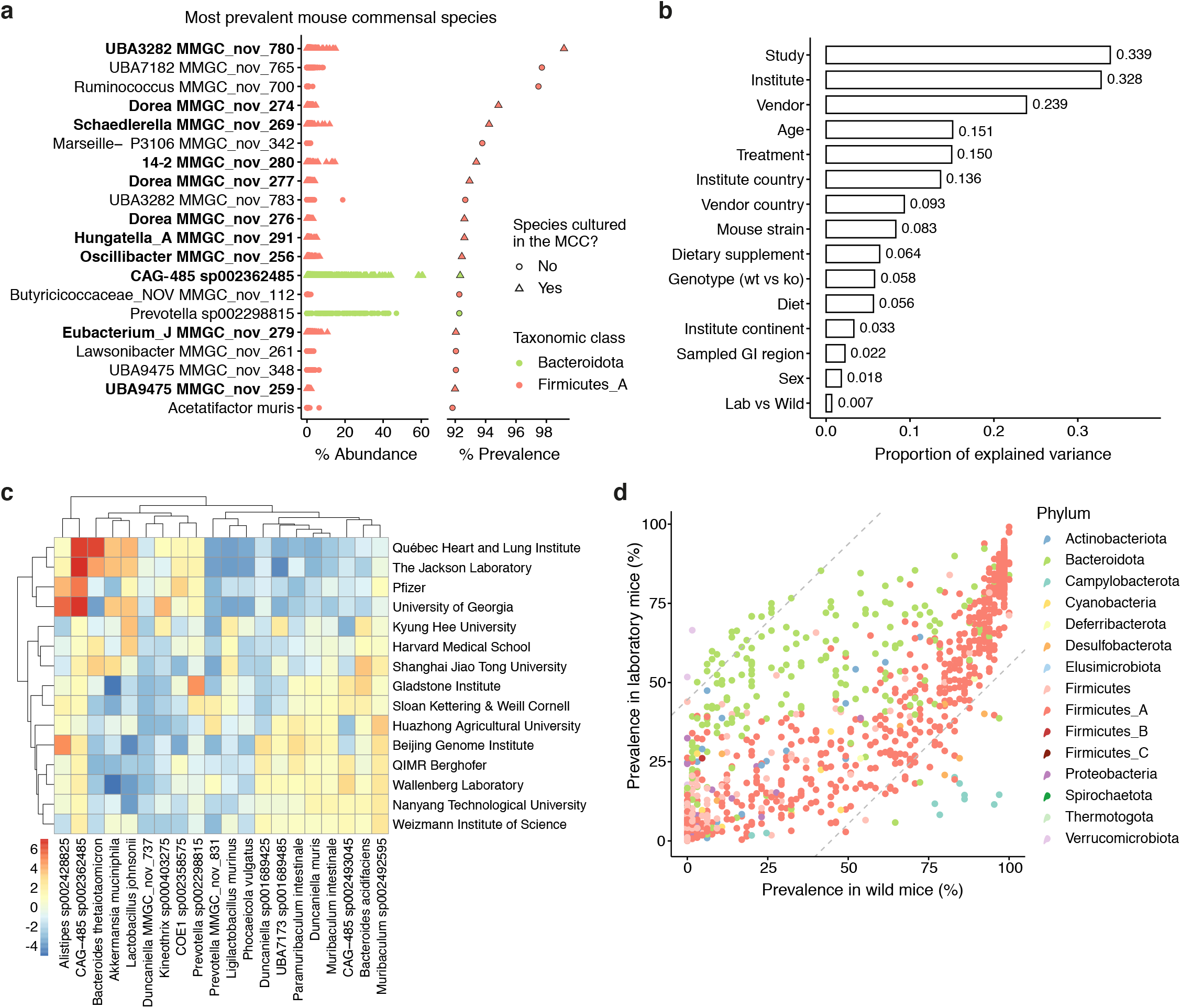
The gut microbiome of the mouse. a) Abundance profiles of the 20 most prevalent species of the mouse microbiota (n=1,924). A species was determined as present in a sample if it was assigned ≥0.01% of classified reads. Point colour represents taxonomic phylum and point shapes and boldface labels indicate whether a species has been cultured as part of the MMGC (triangles) or not (circles). b) Proportion of variance explained (R2) by variables in the metadata using a PERMANOVA test. All analyses were run with 999 permutations. c) Heatmap showing abundance of the top 20 most abundant species of the mouse microbiota across different institutes. Analyses include faecal samples from wildtype C57BL/6 “control” mice fed chow diets. Data are centre log-ratio normalised read fractions, following Bayesian-multiplicative replacement of count zeros. d) Scatter plot comparing prevalence of species between laboratory and wild mice gut microbiomes. Colours represent taxonomic phylum. Dashed grey lines represent two standard deviations from ΔPrevalence(lab vs wild) = 0.

Differences in the host microbiota have been increasingly recognised to be a potential source of irreproducibility in mouse models^24, 25^. We therefore sought to assess the effect of host genetic and environmental factors (vendor, institute, study, genotype, laboratory-versus-wild, diet) on compositional variation of the mouse gut microbiome (Figure 2b). The study itself was the most impactful factor, explaining 33.9% of the variance in microbial community (Figure 2b, permutational analysis of variance, P<0.001, 999 permutations, Supplementary Table 10), followed closely by the institute in which the study was performed (32.8%). To further explore the microbial differences between mice from different institutes, we compared abundance of the 20 most dominant species of the mouse microbiome across 375 C57BL/6 “control” mice from 15 institutes (Figure 2c). Institutes differed greatly in abundance of key microbial species. Phenotypically important bacterial species such as *Bacteroides thetaiotamicron, Lactobacillus johnsonii, Ligilactobacillus murinus* (formerly *L. murinus*) and *Akkermansia muciniphila* were all found to be differentially abundant between institutes. These differences may underpin the irreproducibility observed between mouse studies performed in different institutes^24, 26, 27^.

Recent studies have indicated that the microbiota of wild mice is highly distinct from laboratory mice, influencing outcomes in models of disease^28^. We looked at the differences between the microbiotas of laboratory and wild mice. Surprisingly, only 0.07% of the variance in microbiome composition could be explained by ‘wild vs laboratory’ metadata (Figure 2b), indicating that wild mouse metagenomes were highly similar to those of laboratory mice. Indeed, 90.5% of commensal species are shared between laboratory and wild mice, and species prevalence between these mouse cohorts correlates strongly (*R* = 0.8, *P* < 2.2×10^16^; Figures 2d & S3a). No species were found to be unique to wild mice (Figure S3b), but notably many species of the phyla Campylobacterota and Desulfobacterota were found to be more prevalent in wild mice (Figure S3c), while *Akkermansia muciniphila* was more prevalent in laboratory mice (Figure S3c) and *Akkermansia muciniphila_A* was unique to the laboratory setting (Figure S3b). Taken together, these analyses indicate that while institutional housing environments represent a substantial source of variation, the microbiota of laboratory mice is more similar to that of wild mice than previously suggested.

### Taxonomic versus functional overlap between humans and mice

As the MMGC is a comprehensive representation of the global mouse microbiome, we next determined the degree to which gut microbial species are shared between humans and mice. Using a core genome taxonomy approach to compare species in the MMGC with human gut bacterial species from the Unified Human Gastrointestinal Genome (UHGG) collection^5^, 3,922 species can be attributed to colonising at least one host (Figure 3a), while only 104 species (2.65%) were found to colonise both humans and mice (Figure 3b; Supplementary Table 6). Of these shared species, 97 (92.4%) have been previously characterised, including known pathogens such as *Clostridioides difficile* and common members of the skin microbiota such as *Staphylococcus aureus* and *S. epidermidis*. Because some previous studies have defined the taxonomic overlap between the human and mouse microbiota by using gene-level analyses^9, 10^, we also examined gene cluster sharing between hosts. Using 90% sequence identity as a threshold for gene sharing at the species level, 3.97% of clusters were shared between hosts (Figure 3c), which is consistent with previous estimates using the same method^9^.

**Figure 3.**
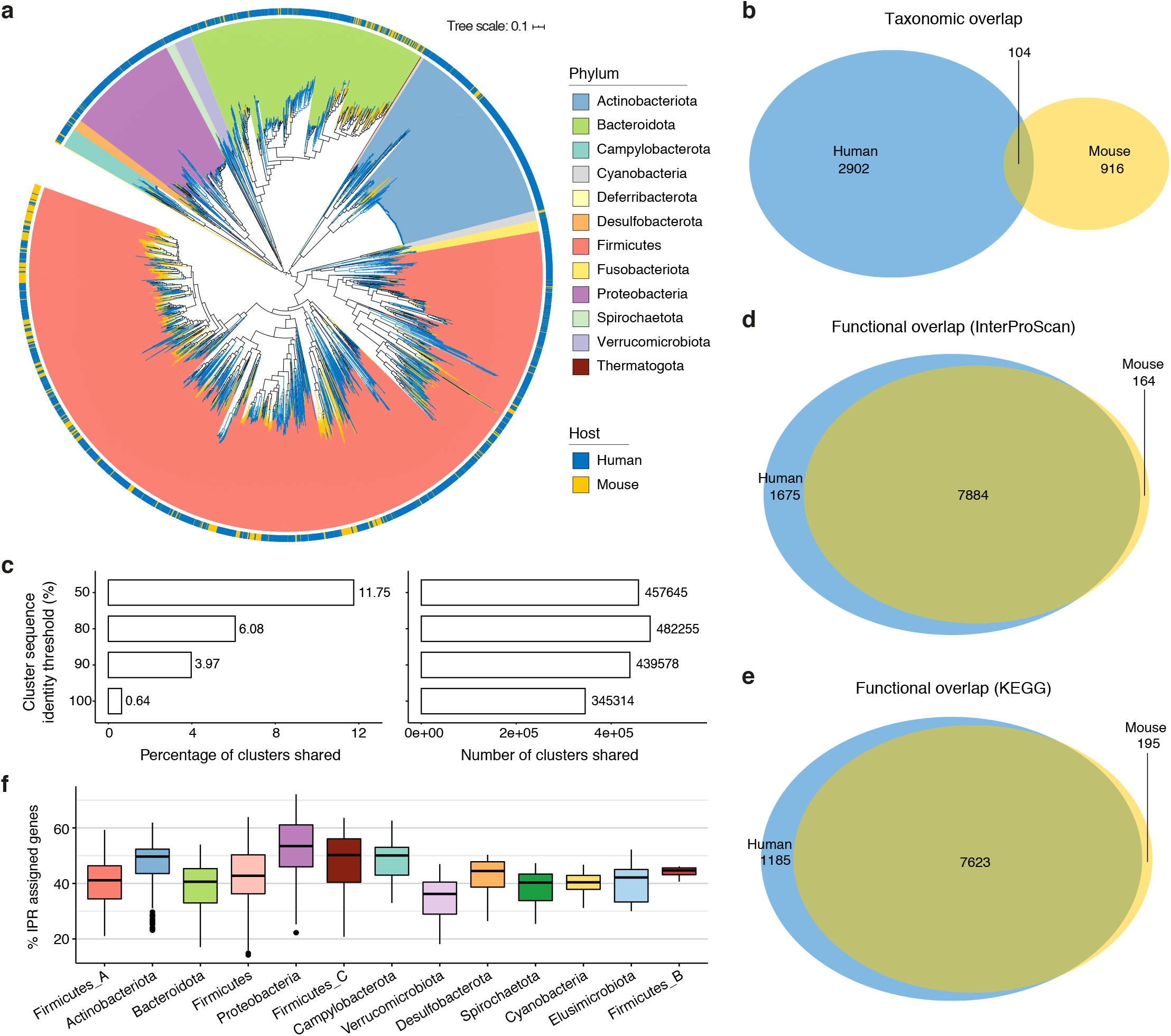
Taxonomic and functional comparisons of species of the human and mouse gut microbiotas. a) Maximum-likelihood phylogenetic tree of 3,006 human and 1,021 mouse representative genomes using 120 core gene alignment. Colour range indicates phylum, the outer colour bar and branch colours indicate host organism. b) Venn diagram illustrating species sharing between human and mouse microbiotas. c) Sharing of gene clusters between hosts. Data represent the percentages (left) and counts (right) of gene clusters that are shared between all human and mouse commensal genomes at different sequence identity thresholds. 90% is an estimate for species level. d-e) Venn diagrams illustrating the functional overlap of InterProScan protein families (IPR; d) or KEGG Orthology (KO) groups (e) between all human and mouse gut commensal species. f) IPR assignment efficiency by taxonomic phylum. Data represent the percentage of predicted protein-coding genes of each pangenome that could be assigned to an IPR protein family, coloured by phylum.

As our approaches indicated that the microbiota of humans and mice were taxonomically very distinct populations, we next sought to explore the degree to which taxonomic overlap corresponded to functional overlap between these populations. To perform these comparisons, we generated pangenomes for each human and mouse commensal species and annotated the predicted protein coding sequences using InterProScan^29, 30^ and eggNOG mapper^31, 32^ to predict pangenome functionality. Of 9,723 InterPro protein families (IPRs) and 9,003 KEGG Orthology (KO) groups^33^ that could be predicted in at least one pangenome, 7,884 IPRs (81.1%; Figure 3d) and 7,623 KOs (84.7%; Figure 3e) were shared in at least one human- and mouse-derived commensal pangenome. Notably, the average proportion of proteins that could be assigned with InterProScan was 42.5% but varied between phyla (Figure 3f). In addition, mouse pangenomes were significantly less annotated (Figure S4a), likely reflecting bias in protein databases for human-derived microbes. Therefore, while we estimate that over 80% of known functions are shared between human and mouse commensals, there are likely many functions which we cannot compare between hosts at this time. Collectively, these analyses show that the mouse and human microbiotas are taxonomically very distinct; however, the functions performed by gut microbes are largely conserved between mouse and human.

### Drug metabolism by the microbiota differs between humans and mice

Given the disparities between taxonomical and functional similarity between microbes from the mouse and human microbiotas, we next sought to identify functionally equivalent species between hosts. The gut microbiota has been increasingly shown to alter drug metabolism, with different commensal species able to activate, inactivate or modify a drug, resulting in different requirements for dosing and increasing the risk of side effects^34, 35^. While mice are commonly used in pre-clinical drug studies for initial dosing and toxicity analyses, the role of inter-host microbiota differences in pharmacology is unknown. To identify potential drug metabolisers within the mouse microbiota, we screened our MMGC for genes that were previously validated for metabolising drugs in the human microbiota. Diltiazem, a commonly prescribed antihypertensive, was recently shown to be metabolised by homologues of the *bt_4096* gene, identified in *Bacteroides thetaiotaomicron* from the human microbiota^36^. In this study, the ability to metabolise diltiazem correlated with *bt_4096* gene abundance rather than the abundance of *B. thetaiotaomicron*. Additionally, *bt_4096* homologues with sequence identity as low as 30% were validated to have the capacity to metabolise this drug^36^. We screened the MMGC for homologues of the *bt_4096* gene and predicted a species to be a diltiazem-metaboliser if more than 50% of its strains encoded a gene with at least 50% sequence identity to *bt_4096*. Whereas nearly all predicted diltiazem-metabolisers in the human host belonged to the genera *Alistipes, Bacteroides* or *Parabacteroides*, a large fraction of predicted mouse diltiazem-metabolisers were found in the *UBA7173* and *RC9* genera, specifically *UBA7173 sp001689485* and *RC9 sp002298075* respectively (Figures 4a). Diltiazem-metabolising species of the human microbiota were also found in the mouse microbiota where they were highly prevalent and relatively abundant (Figure 4b), suggesting that these species may mediate diltiazem metabolism across host microbiotas. In contrast, *Enterococcus faecalis* tyrosine decarboxylase (TyrDC) has been shown to metabolise L-DOPA to dopamine, reducing efficacy in Parkinson’s Disease; the abundance of *E. faecalis* in human faeces strongly correlates with faecal L-DOPA metabolism^37^. Although *E. faecalis* is also found in the mouse microbiota, it is a relatively minor component, with a prevalence of 15.6% and a mean abundance of 0.05% of classified reads. Instead, orthologous TyrDC genes are most commonly found in the *Lactobacillaceae* family in the mouse microbiota (Figures 4c,d). Therefore, although the capacity to metabolise certain drugs is likely conserved between the human and mouse microbiotas, the taxonomic locations of these functions may be largely distinct between hosts.

**Figure 4.**
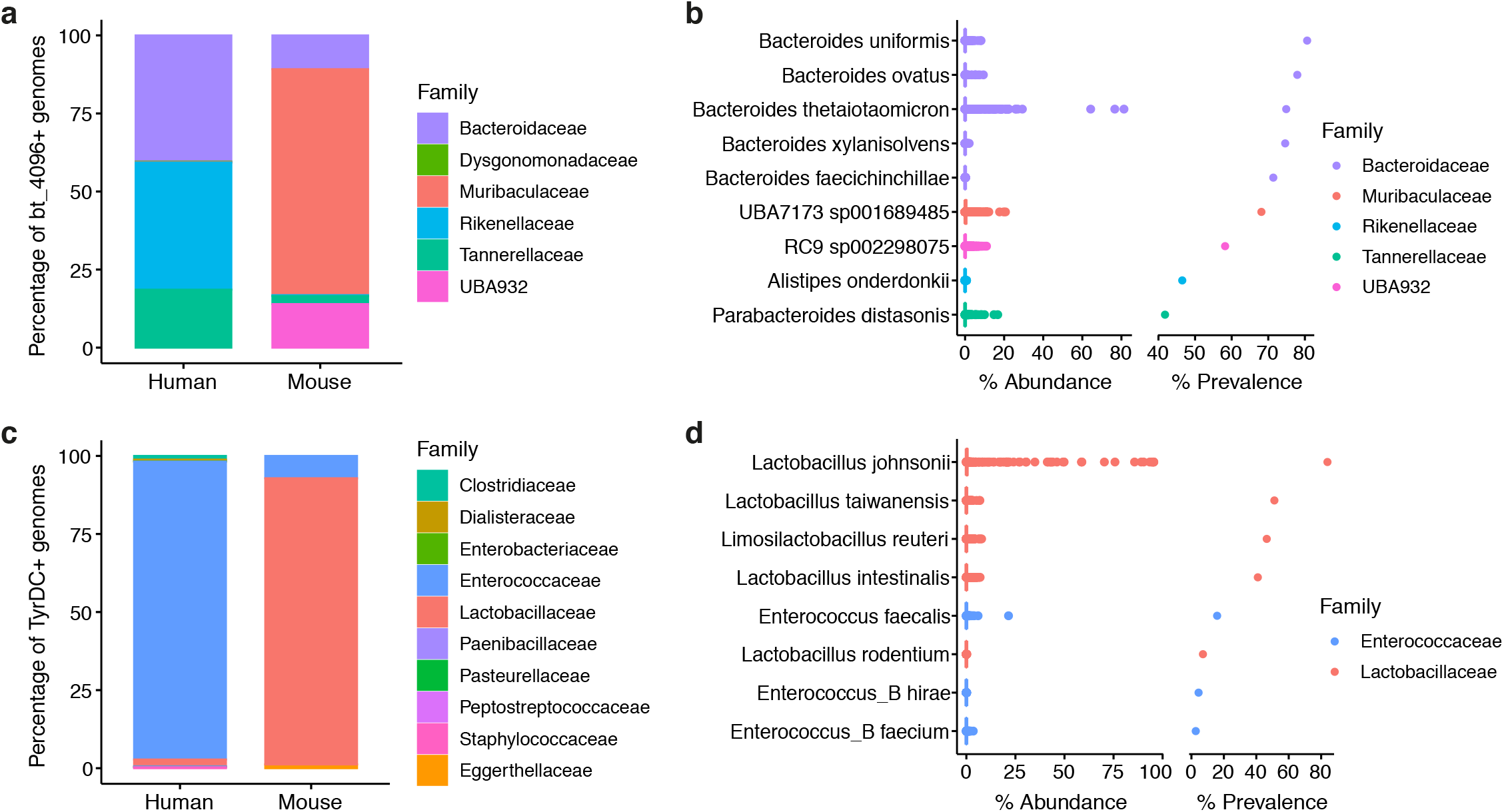
Examples of inter-host drug metabolism by the gut microbiota. a-b) Predicted diltiazem metabolisers encoding the *bt_4096* gene. c-d) Predicted L-DOPA metabolisers encoding the *ef_E6I994* tyrosine decarboxylase. a,c) Stacked bar charts representing the family level taxonomic distribution of *bt_4096* (a) and *ef_E6I994* (c) genes across the UHGG and MMGC. b,d) Percentage read assignment (% abundance) and prevalence (% prevalence) of *bt_4096* (b) and *ef_E6I994* (d) gene-encoding species in the mouse microbiota (n=1,924). A species is determined as present in a sample if it is assigned ≥0.01% of classified reads. Colours represent taxonomic family.

Digoxin is an important heart failure medication that needs to be dosed precisely to avoid side effects. Its metabolism by the human microbiota has been attributed to the *Eggerthella lenta* cardiac glycoside reductase (cgr) operon^38, 39^. In particular, the *Eggerthella lenta* cgr2 gene was shown to be necessary and sufficient for digoxin reduction, while being prevalent in the human microbiota^40^. *Eggerthella lenta* is not found in the mouse microbiota so we explored whether homologues of the cgr2 gene exist in a mouse commensal species that might be functionally equivalent. In humans, a homologous gene product is found in the closely related species *Aldercreutzia equolifaciens*; however, we were unable to find any gene homologue in a mouse commensal species. Likewise, *Dorea scindens* (formerly *Clostridium scindens*) has been shown to encode a 20a-hydroxysteroid dehydrogenase, desC, that metabolises glucocorticoids such as cortisol and dexamethasone^41^. *Dorea scindens* is not found in the mouse microbiota, so we screened the MMGC for desC homologues. No genes with sequence identity above 50% could be found. These therefore represent examples where an important function of the human microbiota may not be reconstituted in the mouse, suggesting potential consequences for the use of mice as models in certain aspects of basic medical and pharmaceutical research.

### Taxonomic locations of butyrate synthetic pathways differ between humans and mice

Some crucial microbial functions have been well-characterised in the human host; however, as a result of the previous scarcity in characterisation of the mouse microbiota the species which encode these functions in the mouse gut have remained unknown. A notable example of this is synthesis of the short-chain fatty acid butyrate, a phenotypically important metabolite that is produced by the microbiota. Butyrate is the main energy source for enterocytes^42^ and is essential for maintaining a healthy, oxygen-free environment in the gut^43, 44^. In addition, it is involved in regulating host metabolism^45^, sleep^46^ and healthy cognitive functioning^47^, it induces peripheral T regulatory cells^48, 49^, and is implicated in diseases as diverse as inflammatory bowel disease^50^ and depression^51^. Butyrate is synthesised from dietary fibre or from amino acids such as glutamate and lysine, all of which culminate in the conversion of butyryl-CoA to butyrate via either butyrate CoA-transferase (BCoAT; direct) or via a phosphotransferase pathway (PTB/BUK; indirect)^52^ (Figure 5a). While the direct pathway is thought to be the main pathway for butyrate synthesis by the microbiota in humans^53, 54^, our analyses revealed that the phosphotransferase pathway appears to be more dominant in the mouse host (Figure 5b), although both pathways are prevalent in the mouse microbiome (Figure 5c).

**Figure 5.**
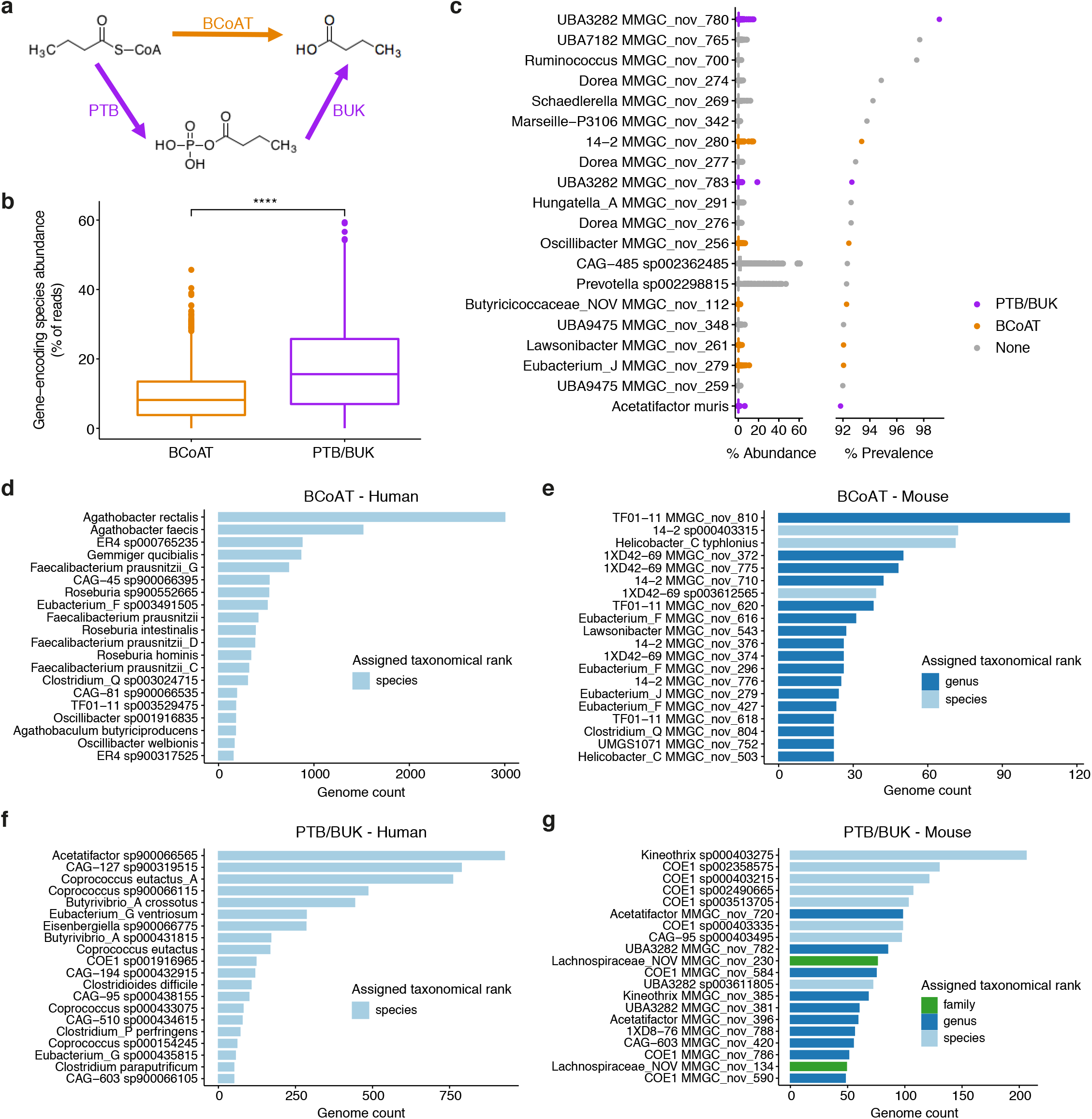
Butyrate metabolism is performed by different species in the human and mouse microbiotas. a) Schematic of the terminal pathways of butyrate synthesis by the gut microbiota. Butyrate CoA-transferase (BCoAT; orange); butyrate phosphotransferase/butyrate kinase (PTB/BUK; purple). b) Pathway abundance estimation using genome-encoding species abundance. A pangenome species was designated as a butyrate-producer if >80% of its contributory genomes encoded the relevant terminal pathway gene. PTB/BUK pathway is the most abundant pathway for butyrate synthesis in the mouse gut microbiota. c) Abundance profiles and predicted butyrate-producing status of the 20 most prevalent species of the mouse microbiota. Colour represents the encoded terminal pathway for butyrate synthesis; no species encoded both pathways. d-g) Bar charts of the top 20 butyrate pathway-encoding species in humans and mice. Data represent counts of gene-encoding genomes per species. d) top BCoAT-encoding species in the UHGG, e) top BCoAT-encoding species in the MMGC, f) top PTB/BUK-encoding species in the UHGG, g) top PTB/BUK-encoding species in the MMGC. Colours represent the lowest taxonomic rank assigned to a species by GTDB-Toolkit.

BCoAT genes can be found in humans in model butyrate producers such as *Agathobacter rectale* (formerly *Eubacterium rectale*), *Faecalibacterium prausnitzii*, and *Anaerostipes hadrus*^50^ (Figure 5d); however, none of these species is represented in the MMGC. Instead, an overwhelming majority of BCoAT genes in the MMGC were found in previously uncharacterised species that are unique to the mouse gut microbiota (Figure 5e), indicating unprecedented novelty in mouse butyrate producing populations. At the genus-level, while the majority of BCoAT genes in human commensals were found in *Agathobacter* and *Faecalibacteria* (Figure S5a), in mice the BCoAT gene was instead found in the *14-2* genus, while a member of genus *TF01-11* has the most representative BCoAT gene-encoders. Therefore, although the BCoAT pathway is conserved between the human and mouse microbiota, the taxonomic location of this metabolic capability differs between hosts.

The second pathway for butyrate synthesis from butyryl-CoA involves butyrate phosphotransferase and butyrate kinase (Figure 5a). This pathway is commonly represented in the human microbiota by species of *Coprococcus*^*52, 55*^ and other species of the *Clostridium* cluster *XIVa*^*52*^ (Figure 5f, S5b). Similar to the BCoAT-encoding species, the functionally equivalent members of the mouse microbiota in the MMGC were largely previously uncharacterised (Figure 5g). This pathway predominated in the *COE1* and *Kineothrix* genera (Figure S5b). Collectively, these interrogations of the mouse microbiota using the MMGC enable previously unachievable comparisons to the human microbiota and demonstrate that conserved metabolic functions may be performed by very different taxa between hosts. In addition, they identify the specific commensal mouse microbes that house these key functions and reveal areas in which further mouse microbiota studies may or may not be highly informative.

## Discussion

In an endeavour to overcome existing gaps in knowledge regarding the utility of mouse microbiota models for the study of human physiology and disease, we set out to better define the global mouse microbiome composition by developing a comprehensive collection of mouse gut bacterial genomes and cultures. Our collection, the MMGC, both improves correlative analyses of mouse metagenomes and facilitates the study of causation in the mouse microbiota through availability of cultured isolates. The MMGC is the largest and most complete collection of mouse commensal genomes to date, allowing us to develop an extensive set of additional tools to perform more accurate and detailed comparisons between the mouse and human microbiotas. These comparisons revealed that while key metabolic functions of the microbiota are conserved between mice and human, they may be performed by distinct identifiable microbes in each host.

While these studies represent a substantial advance in accessing and understanding the role of the microbiota in biomedical research, a number of limitations remain. Although our culturing methods have yielded 276 isolates from 130 species, approximately 87% of species remain to be cultured, and this must be prioritised for future studies. To analyse MAGs and isolate genomes together, we used stringent quality criteria to ensure that only the most reliable genomes were considered. One drawback of this approach, however, is that species that cannot be given accurate levels of completeness of contamination by current tools may be excluded from analyses. Nevertheless, accurate functional analyses required a consistent level of genome completeness to make analyses between genomes comparable. In addition, while predicted functionality is largely conserved between the human and mouse microbiotas, facilitating the use of mouse models for microbiota-susceptible phenotypes, it is important to consider that 20% of functionality is not conserved between hosts. Furthermore, our functional analyses are only as complete as the functional annotation of bacterial genes, leaving roughly half of the functional potential of the average genome unknown. The MMGC can therefore best be used to identify if and in which species a known function of interest is conserved between the mouse and human microbiotas to determine if a suitable mouse model can be developed.

Irreproducibility between mouse models is a major concern in biomedical research, and this is hypothesised to be at least in part due to differences in the microbiota. The improved metagenomic analyses enabled by the MMGC now allow us to begin to address these challenges by facilitating the accurate quantification and benchmarking of the mouse microbiome. Mice and humans have co-evolved with their respective microbiotas, and given the extent of taxonomic divergence as well as potential functional dichotomy, the implications of inter-host microbiota transplantations are currently unknown. Studying commensals in their respective hosts is likely to be the most physiologically valid approach, and this work provides the ability to define the closest functionally equivalent species for translational research between mice and humans.

## Methods

### Bacterial culturing

Fresh faeces were collected from 30 different mice with a range of genotypes, across 10 different mouse colonies at the Wellcome Sanger Institute (Supplementary Table 7). Faeces were homogenised in sterile PBS (100mg/mL) and the faeces mashed up and vortexed. A 1 in 10 dilution series was done into sterile PBS and 200µL of each dilution plated onto a range of large agar plates (Supplementary Table 8) and incubated at 37C. This was done aerobically and anaerobically onto pre-reduced plates in a Don Whitley anaerobic workstation (80% nitrogen, 10% carbon dioxide, 10% hydrogen). After 2 days, individual colonies were picked onto fresh agar plates and left to grow, single colonies were then picked again onto fresh plates to ensure purity. A colony prep was carried out on each isolate by scraping a loop full of growth into a 2ml screw cap tube containing glass beads (acid-washed 425-600µm) and 500µl sterile PBS. This was placed into a MPBio FastPrep Instrument and shaken at speed 6.0 for 30 seconds. After centrifugation at 14000rpm for 5 minutes, 1µL of the supernatant was used to carry out a 16S PCR using the standard 7F (AGAGTTTGATYMTGGCTCAG) and 1510R (CCTTCYGCAGGTTCACCTAC) bacterial primers and Promega GoTaq Hot Start reagents. Any PCR products were sequenced and MOTHUR used to align the resulting sequences and create OTUs. A sequence from each OTU was identified by NCBI BLAST and an isolate selected from each new OTU identified. 10mL of BHI or YCFA broth was inoculated for each new isolate identified and left to grow overnight. 500µL of the overnight culture was mixed with 500µL of 50% glycerol in a cryotube (performed in quadruplicate) and these were frozen at -80C. The remaining overnight culture was spun at 4000rpm to create a cell pellet and then washed in PBS. The washed pellet was then used to extract genomic DNA using the ‘Lucigen MasterPure Complete DNA and RNA Purification Kit’. Genomic DNA was kept at 4C until used for whole-genome sequencing on a HiSeq X 10, 150bp PE, 192 plex.

### Genome sequencing and assembly

Genomic DNA was sequenced using the Illumina Hi-Seq Ten platform at the Wellcome Sanger Institute with library fragment sizes of 200–300 bp, a read length of 150 bp and a target read depth of 100x. Annotated assemblies were produced using the pipeline described previously^56^. For each sample, sequence reads were used to create multiple assemblies using Velvet v1.2^57^ and VelvetOptimiser v2.2.5^58^. An assembly improvement step was applied to the assembly with the best N50, and contigs were scaffolded using SSPACE^59^ and sequence gaps filled using GapFiller^60^. Automated annotation was performed using PROKKA v1.11^61^. A total of 288 cultured isolates were submitted for whole genome sequencing, however only 276 passed our QC checks (see below) and were used in our analyses.

### Public metagenomes and genomes

We curated 1,960 publicly available mouse metagenomes from NCBI (last accessed in June 2020) and MG-RAST (last accessed in February 2020). Only samples from mice that had not received treatments that could disrupt the endogenous status of their microbiota, such as antibiotic therapy and human faecal microbiota transplantation, were curated for this study.

In addition, NCBI was searched for mouse gut-derived bacterial genomes from previous studies (last accessed March 2020), including the miBC^6^ and mGMB^8^, as well as other smaller collections of genomes with mouse gut metadata. 330 public isolate genomes were curated, but only 274 passed our stringent quality control criteria.

The Unified Human Gastrointestinal Genome^5^ (UHGG) v1.0 collection was used as the sole source for human gut-derived bacterial genomes. 204,939 non-redundant genomes were downloaded and subjected to the same quality control and taxonomy assignment pipeline as the genomes of the MMGC. In total, 100,456 non-redundant, near-complete human gut microbial genomes, representing 3,006 species (Supplementary Table 9), were curated and compared to genomes of the MMGC.

### MAG synthesis

MAGs were generated using a custom in-house pipeline that leveraged MetaWRAP v1.2.2^62^ to assemble, bin and refine genome bins. Metagenomes were initially quality controlled using KneadData v0.7.3 with default settings. Host reads were removed from samples using the GRCm39 reference genome and Bowtie2 v2.3.5^63^. In addition, reads were aligned to the Phi X 174 genome and removed. For single-sample assembly, paired-end samples were assembled using metaSPAdes v3.10.1^64^ and single-end samples, as well as paired-end samples that failed to be assembled with metaSPAdes, were assembled with megahit v1.1.1-2-g02102e1^65^. We chose a hybrid binning approach to generate genome bins from assembled contigs as this approach combines the strengths of multiple binning tools to produce higher quality MAGs that are more complete than using binners such as MetaBAT2^62^ in isolation. MetBAT2 v2.9.1^66^, MaxBin2 v2.2.4^67^ and CONCOCT v0.4.0^68^ were used in parallel to produce genome bins that were then consolidated and refined.

### Genome quality assessment

We used CheckM^69^ to define the completeness and contamination of isolate genomes and MAGs. We defined medium-plus quality MAGs as >50% completeness, <5% contamination and a quality score ≥50, where QS = Completeness – (5 * Contamination)^22^. These genomes thereby exceed the medium quality thresholds defined by MIMAG^21^. We generated 45,218 medium-plus quality MAGs as part of this study. For our analyses, we only included genomes that met additional quality criteria: >90% completeness; <5% contamination; maximum genome size ≤ 8 Mb^70^; maximum contig count ≤ 500^12^; N50 ≥ 10 kb^15^; mean contig length ≥ 5 kb^12^. These stringent criteria ensure that only the most complete genomes with minimal fragmentation are compared. These genomes are defined as near-complete^16, 22^.

### Taxonomic assignment and representative genomes

To remove redundancy from our collection of near complete genomes, we used dRep v2.5.4^71^ (--pa 0.999 --SkipSecondary) to remove conspecific genomes that shared ≥99.9% ANI. We used the Genome Taxonomy Database-Toolkit (GTDB-Tk) v1.3 ‘classify_wf’ workflow to assign taxonomic ranks to our genomes according to the GTDB r95 taxonomy. Genomes that could not be assigned a species-level were considered “previously uncharacterised”. Using this approach, 98,841 (98.4%) human and 8,566 (47.4%) mouse gut bacterial genomes were assigned a species-level taxonomy. Novel genomes were dereplicated with a two-step genomic distance analysis using dRep v2.5.4 (-comp 50 -con 5 -pa 0.9 -sa 0.95 -nc 0.6). Previously calculated quality data were supplied for each genome with the --genomeInfo flag to speed up analyses. Genomes that shared ≥95% ANI at 0.6 alignment fraction were considered the same species^72-74^. To determine genome representatives for known species, we ranked each genome according to a quality score,

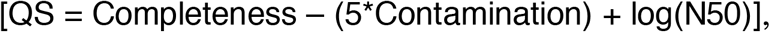

and used the highest scoring genome from each species as the representative.

Although the scope of our study was limited to bacterial genomes only, we looked for archaeal bins that had been generated by our pipeline, however none had been reconstructed from any mouse metagenome.

### Phylogenetic tree building

Protein sequence alignments of 120 core genes generated by the GTDB-Tk ‘align’ module were converted to maximum-likelihood phylogenetic trees using FastTree v2.1.10^75^ with default settings (BLOSUM45 matrix; JTT+CAT model). Trees were visualised using Interactive Tree Of Life (iTOL) v5.6.3^76^.

### Metagenome analyses

Metagenome taxonomic classification was performed using Kraken2^77^ and Bracken^78^. To query the ability of the MMGC to improve metagenomic analyses, we built custom Kraken2 databases for previous mouse commensal culturing studies (miBC, mGMB), all public genomes (Public), the isolates cultured in this study (MCC), representative genomes for the MMGC and a human database containing representative genomes from the UHGG. Only metagenomes that were sufficient read depth to generate MAGs were used in analyses. 1,259 metagenome samples were analysed in parallel with the different Kraken2 databases, and percentage read classification was utilised as a proxy for database efficiency.

To analyse the global mouse microbiome, we used Kraken2 and Bracken custom databases built from MMGC representative genomes. Resulting Bracken outputs were compiled and analysed in R v4.0.2 to determine the most abundant and prevalent species of the mouse microbiome. Due to the compositional nature of metagenome analyses, we determined the Aitchison distances between samples^79^. We performed Bayesian-multiplicative treatment of count zeros^80^ using the zCompositions v1.3.4 R package^81^ and transformed data using a centre log-ratio transformation. Finally, the Euclidean distances of samples were determined using the vegan v2.5-6 package^82^. To assess the ability of metadata variables to explain variance in microbial communities of the mouse microbiome, the adonis function (vegan) was used to run Permutational Multivariate Analysis of Variance on the Aitchison distance matrices using 999 permutations (Supplementary Table 10).

For institute analyses, samples from “control” C57BL/6 mice were curated that were 1) faecal samples, 2) from wildtype mice, 3) not exposed to a wild mouse microbiota, 4) fed only a normal chow diet. Centre log-ratio transformation of the data were performed and a heatmap generated using the pheatmap v1.0.12 package for the top 20 most abundant species of the average mouse microbiome against institute.

### Taxonomic comparisons

We utilised two complementary approaches to define the taxonomic overlap between human and mouse microbiotas: whole genome comparison and gene clustering. For taxonomic comparison, species were considered shared between humans and mice if they were identified as the same species by GTDB-Tk, or, if they could not be assigned at species level, the representative genomes shared ≥95% ANI. For gene-clustering, we concatenated 76,937,350 pre-clustered human predicted proteins for non-redundant, near-complete genomes of the UHGG^83^ with 45,598,646 mouse predicted protein-coding sequences from non-redundant, near-complete species of the MMGC, and performed protein clustering using the ‘linclust’ function^84^ from MMseqs2^85^ v10-6d92c (-c 0.8 --cov-mode 1 --cluster-mode 2 -- kmer-per-seq 80). Proteins were clustered at 100%, 90%, 80% and 50% sequence identity; clusters were considered shared if they contained genes from both human and mouse commensals.

### Pangenome synthesis and functional annotation

To build pangenomes for each species, we concatenated protein-coding sequences from all member genomes and functionally annotated pangenomes using both InterProScan v5.39-77.0^29, 30^ and eggNOG emapper v2.0.1^31, 32^. Functional overlap was determined as the number of IPR protein families and KO groups that were shared between human and mouse commensals.

### Predicting drug metabolism in species of the microbiota

Using well-validated drug metabolism genes from published literature, we used blast v2.7.1 to determine the presence of homologues of these genes in our genome collections. A genome was considered to encode the gene if a blast homologue ≥50% identity alignment was present. If >50% of genomes of a species encoded a drug metabolism function, then that species was considered a drug metaboliser.

### Butyrate synthesis analyses

The InterProScan database was used to locate the terminal pathways of butyrate synthesis in genomes, using the IPR family identifiers: IPR023990 (BCoAT), IPR011245 (BUK), IPR014079 (PTB). Only species encoding both BUK and PTB were considered as butyrate producers using the BUK/PTB terminal pathway. We additionally generated analyses with eggNOG, but the resulting KEGG annotations missed a large number of BCoAT genes, for example in model butyrate producers such as *Faecalibacterium prausnitzii*, so InterPro was deemed the more appropriate database for these analyses.

## Data availability

Raw sequencing data for the isolates of the MCC have been deposited in the ENA under project PRJEB18589. Genome assemblies of the non-redundant, near-complete MAGs and the genome assemblies for the MCC isolates will be deposited under project PRJEB41512 and are currently available for download on the ENA FTP server (http://ftp.ebi.ac.uk/pub/databases/metagenomics/genome_sets/mmgc/). Bacterial isolates will be deposited at the Leibniz Institute DSMZ-German Collection of Microorganims and Cell Cultures (http://www.dsmz.de).

## Acknowledgements

V.A.P. is supported by a Sir Henry Dale Fellowship jointly funded by the Wellcome Trust and the Royal Society [206245/Z/17/Z]. B.S.B. is supported by a studentship from the Rosetrees Trust [A2194].

## Author Contributions

B.S.B., T.D.L. and V.A.P. conceived the study. S.F. initiated culturing of isolates. M.D.S., G.N., E.V., H.P.B. and B.S.B. cultured isolates. B.S.B. curated genomes and metagenomes, generated MAGs, performed computational analyses and analysed and assembled data. N.K., K.V. and A.A. advised on computational analyses. A.A. and B.S.B. wrote code for computational analyses. B.S.B., T.D.L. and V.A.P. wrote the paper.

---

**Supplementary Figure 1:**
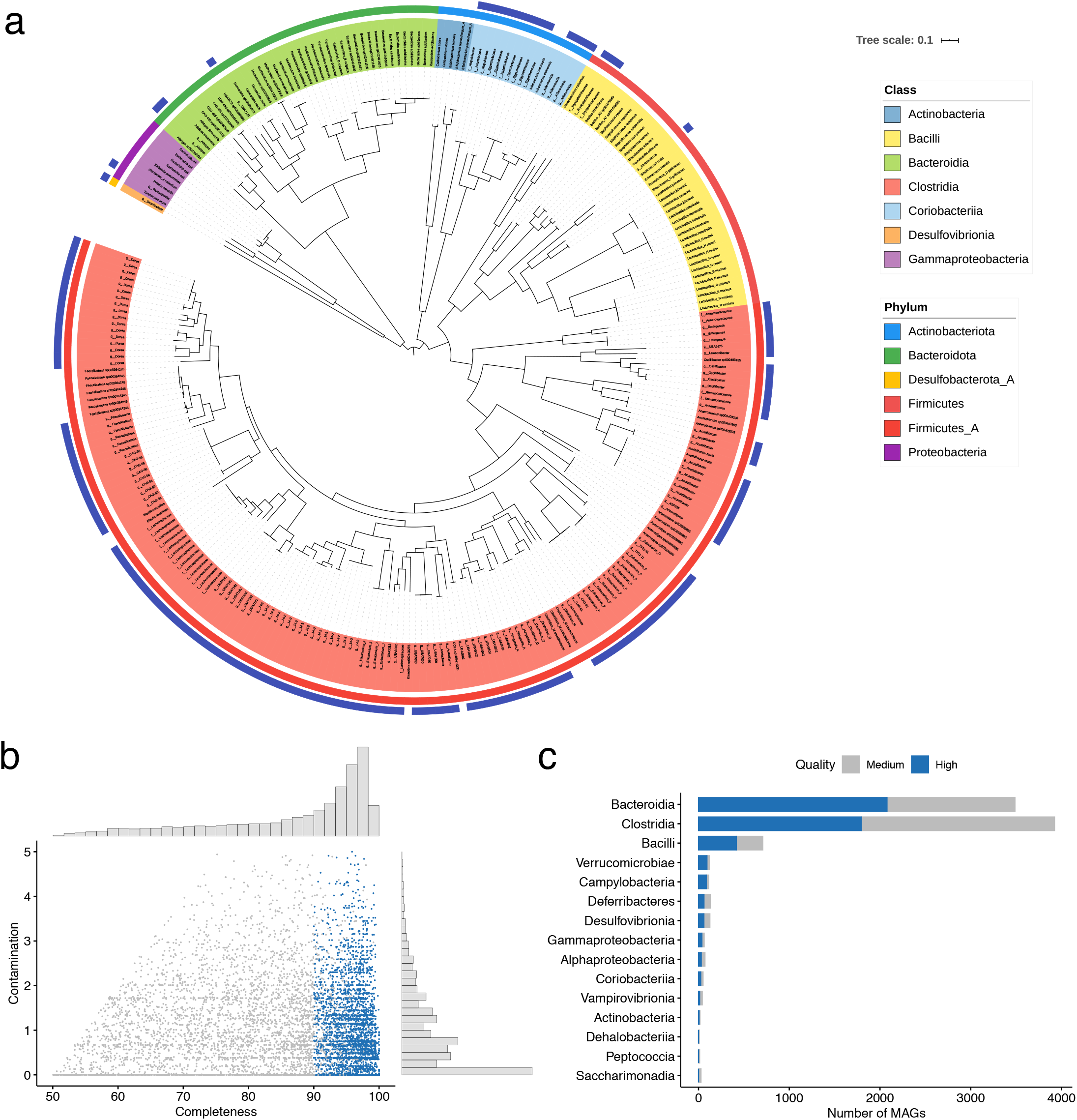
Isolates and MAGs of the MMGC. a) Maximum-likelihood phylogenetic tree of 276 mouse gut bacteria isolated from SPF faeces as part of this study, termed the Mouse Culture Collection (MCC). Tree distances represent alignments of 120 core genes. Labels are coloured according to taxonomic class, while the inner ring indicates GTDB phylum. The outer ring indicates genomes which could not be classified at the species level by GTDB-Tk; labels for these genomes indicate lowest taxonomic rank and taxon. b) Completeness and contamination as determined by CheckM for medium-plus quality MAGs. Blue data points are high quality MAGs, and grey medium. c) Representative cohort of medium-plus MAGs generated in this study, stratified by taxonomic class. Stacked bars show the proportion of high and medium quality MAGs.

**Supplementary Figure 2:**
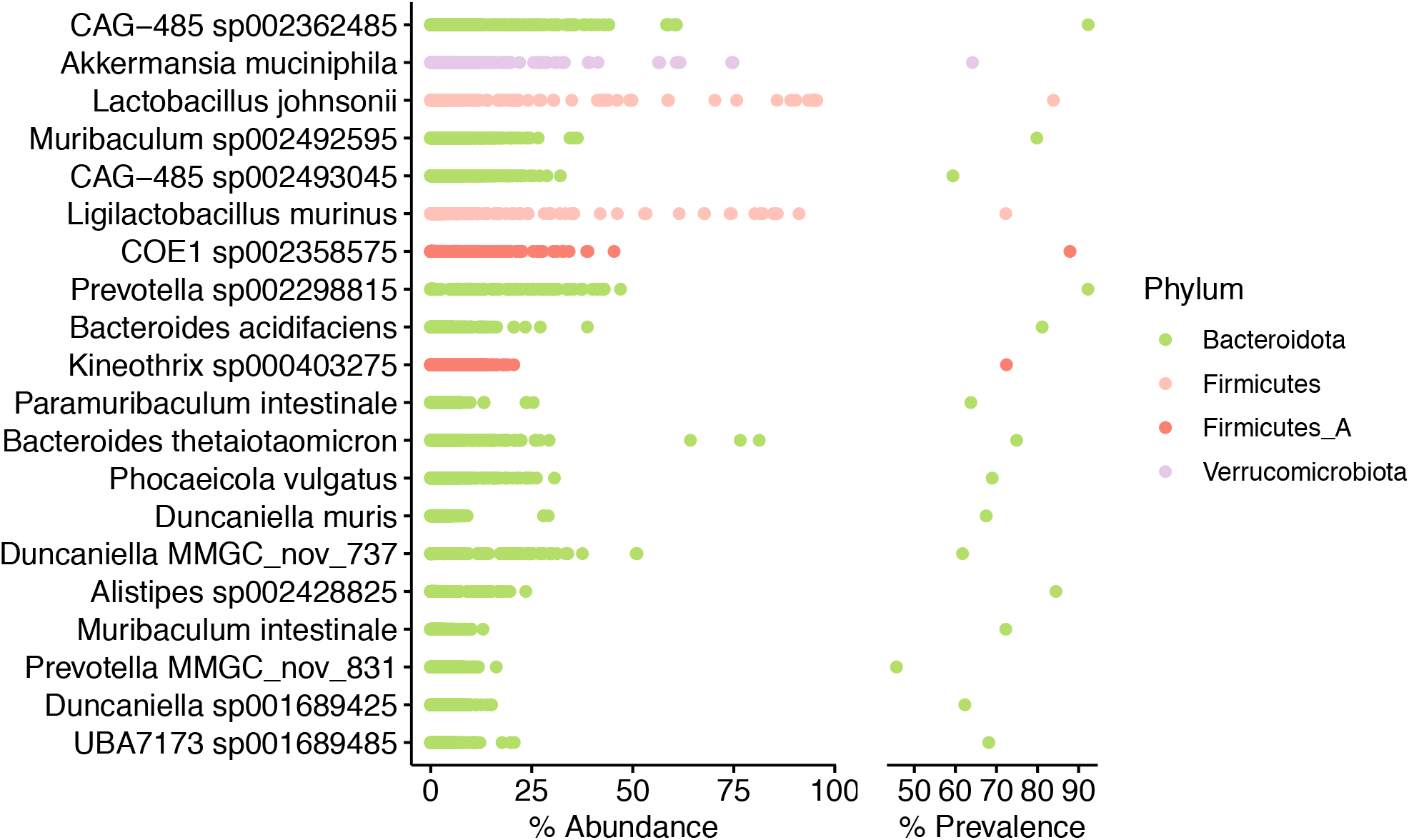
Exploring dominant features of the mouse microbiota. Top 20 most abundant species of the mouse microbiota (n=1,785). Colours represent taxonomic phyla.

**Supplementary Figure 3:**
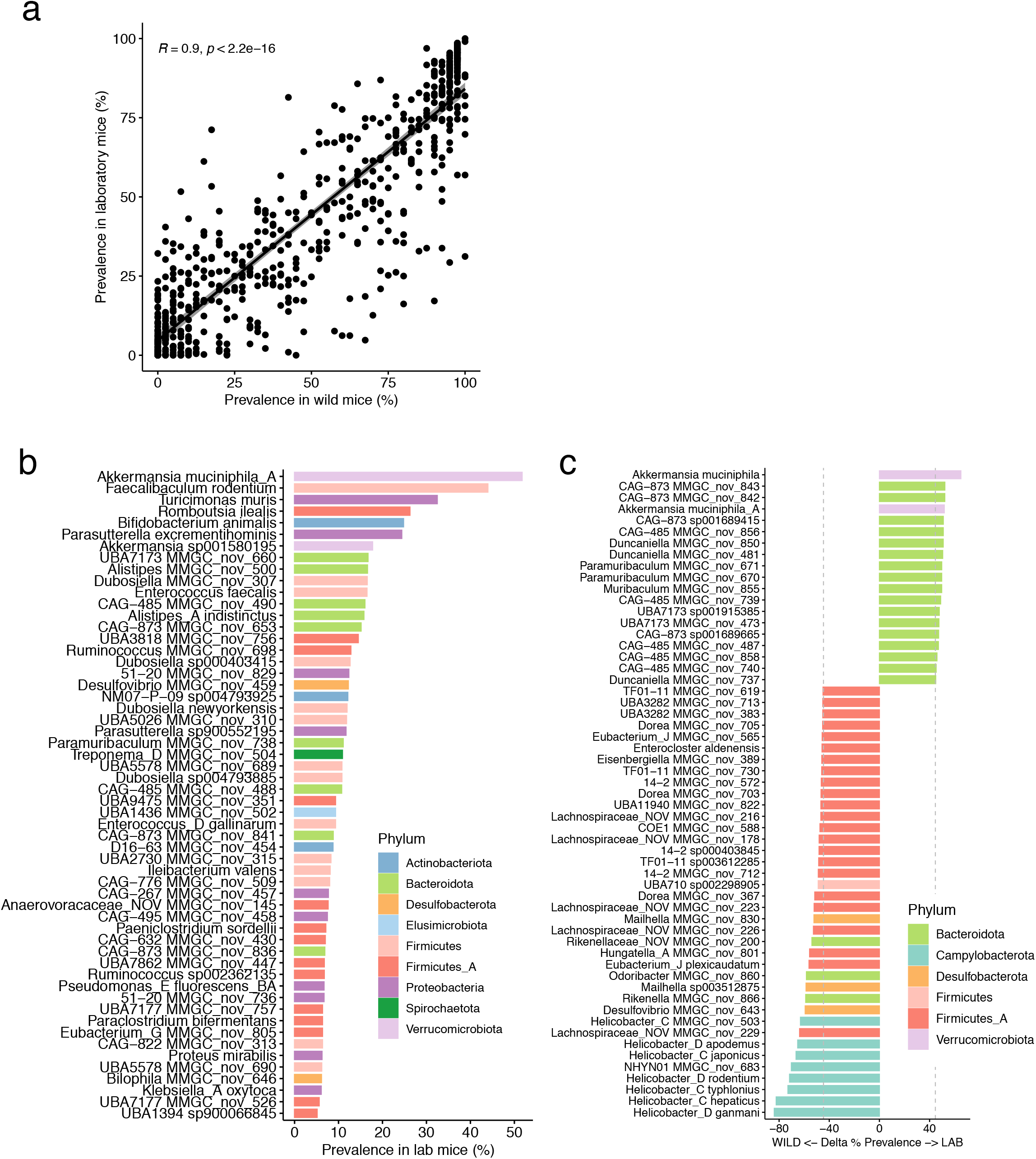
Laboratory and wild mice have similar microbiotas. a) Prevalence of gut metagenome species correlates positively and significantly between wild and laboratory mice. b) Species unique to the laboratory mouse microbiota. Data are prevalence in laboratory mice, colours represent phyla. c) Differential prevalence of gut commensal species between wild and laboratory mice. Only species with a delta prevalence ≥2 standard deviations from 0 (grey dashed lines) are shown. Colours represent phyla.

**Supplementary Figure 4:**
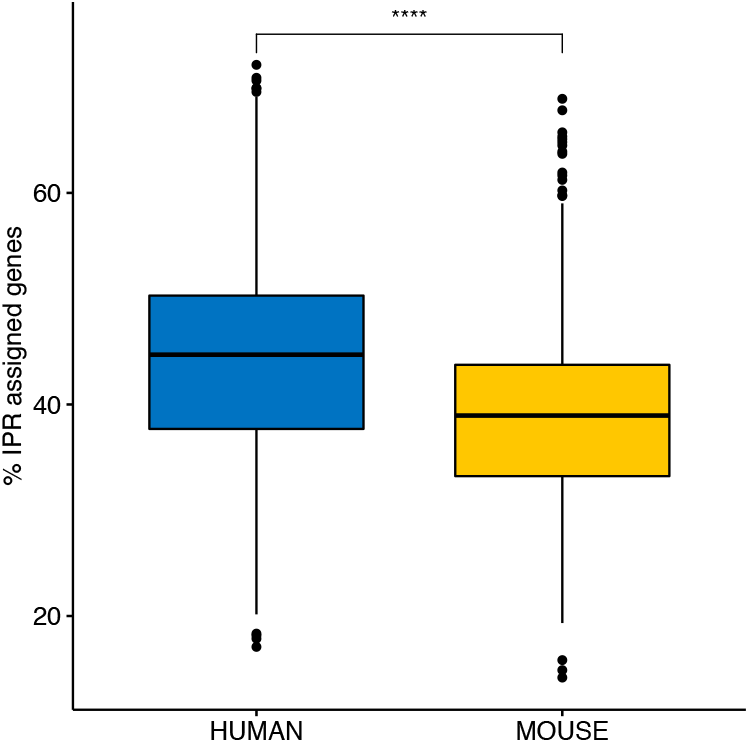
Functional databases demonstrate bias between human and mouse pangenomic species. Box plots showing percentage of predicted pangenome protein-coding sequences annotated with InterProScan protein families (IPRs).

**Supplementary Figure 5:**
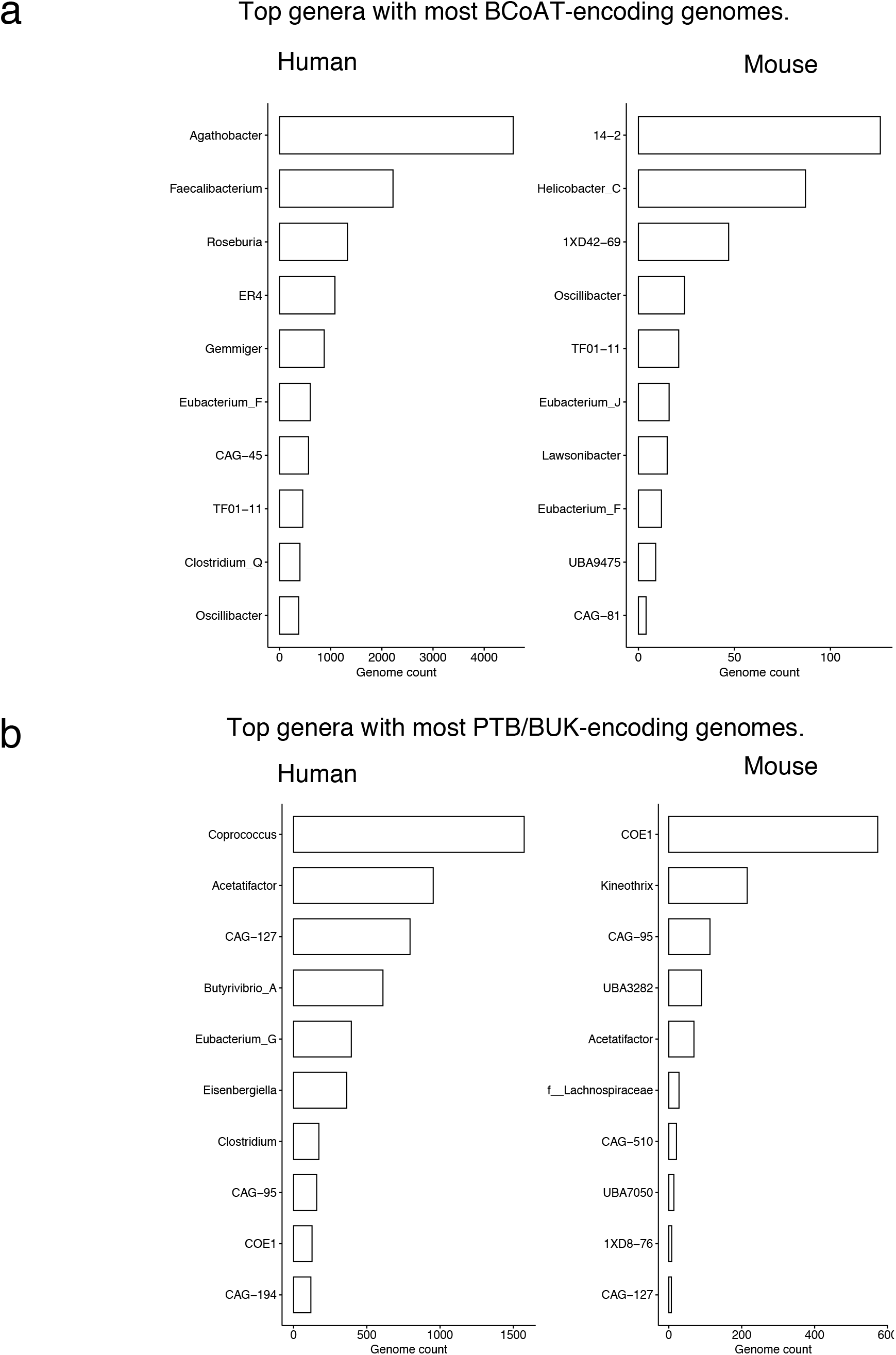
Butyrate-producing species of the mouse and human microbiotas diverge at the genus level. a) Top butyrate CoA-transferase (BCoAT)-encoding genera and (b) butyrate phosphotransferase/butyrate kinase (PTB/BUK)-encoding genera in the UGHH (left) and MCGC (right). Data represent counts of genomes that encode genes for the relevant terminal pathway.

